# MyeliMetric: A Python-Based Toolbox for Standardized G-ratio Analysis of Axon-Myelin Integrity

**DOI:** 10.1101/2025.07.14.662606

**Authors:** Intakhar Ahmad, Farjana Sultana Chowdhury, Anne I. Boullerne, Alexander Gow, Douglas L. Feinstein

## Abstract

The g-ratio, defined as the ratio of an axon’s diameter to the total fiber diameter (axon plus myelin), is a key metric for assessing myelin integrity and axonal conduction velocity in both the central and peripheral nervous systems. Deviations from the physiological range often signal underlying pathology. Despite its diagnostic importance, there is currently no standardized, open-source tool for g-ratio analysis from post-segmented electron microscopy images. To address this gap, we developed MyeliMetric, a Python-based, user-friendly toolbox that streamlines g-ratio data preprocessing and integrates biologically informed validation, requiring minimal statistical expertise to operate without introducing common analytical errors. It is built on the principle that g-ratios exhibit relative consistency across varying axon diameters in healthy conditions. To rigorously assess this relationship, MyeliMetric implements a binning strategy that groups axons into biologically relevant diameter cohorts, enabling the detection of size-dependent deviations in g-ratio distributions. This approach addresses common limitations in conventional analyses, including insufficient sampling, pseudo-replication, and artifacts such as misleading regression slopes. Validation using both synthetic and published datasets from rodent models of demyelination demonstrated the tool’s accuracy, reproducibility, and biological relevance. Synthetic data yielded expected outcomes, and in experimental models, MyeliMetric reliably detected reductions in myelin thickness through g-ratio shifts while minimizing artifacts, thereby providing biologically meaningful insights. It is available on GitHub: https://github.com/Intakhar-Ahmad/NeuroMyelin-G-Ratio-Analysis-Toolkit

## Introduction

Myelin sheaths are essential components of the central and peripheral nervous systems, enabling saltatory conduction that accelerates signal transmission along axons while reducing conduction time and metabolic demand (Nave and Werner 2014; Bechler and Ffrench-Constant 2014). Proper myelination is fundamental to cognitive, sensory, and motor function, as well-myelinated axons transmit action potentials more rapidly and reliably, supporting core neural processes (Stadelmann et al. 2019; Fields 2008). A key quantitative parameter for assessing myelin integrity is the g-ratio, defined as the ratio of the inner axonal diameter to the total outer fiber diameter (axon plus myelin), originally proposed to optimize conduction velocity (Rushton 1951) and later adapted for in vivo imaging applications using MRI techniques (Campbell et al. 2018).

In mammals, optimal g-ratios typically fall between 0.6 and 0.8, reflecting a physiological balance between conduction velocity, structural constraints, and energetic efficiency (Chomiak and Hu 2009; Stikov et al. 2015; Rushton 1951). Deviations from this range are observed in demyelinating conditions such as multiple sclerosis (MS), hereditary neuropathies like Charcot-Marie-Tooth disease, and in experimental models including experimental autoimmune encephalomyelitis (EAE) and cuprizone-induced demyelination (Dupree et al. 2015; Stadelmann et al. 2019; Nave and Werner 2014). These shifts often signal impaired neuronal function, emphasizing the need for precise and reproducible g-ratio quantification.

Despite the g-ratio’s diagnostic and research utility, its broader application is hindered by methodological inconsistencies (Gow 2025). While segmentation tools like AxonDeepSeg (Zaimi et al. 2018) and MyelTracer (Kaiser et al. 2021) delineate axons and myelin from electron or light microscopy images, they lack integrated support for standardized post-segmentation analysis. Without a unified processing pipeline, researchers frequently resort to ad hoc methods such as manual data handling in Excel or custom-written scripts. These practices introduce variability dependent on the user’s skill level, compromise reproducibility, and heighten the risk of analytical errors. As a result, this methodological gap can lead to inconsistent statistical practices and fragmented results across different studies.

A major pitfall in many studies is interpreting results based on analysis of the relationship between g-ratio and axon diameter. However, recent studies by Gow and colleagues (Gow 2025; Gow et al. 2025) have re-visited the importance of analyzing g-ratio data within the context of the axomyelin unit model, a biologically-grounded framework which asserts a linear relationship between axon and fiber diameters, consistent with classical findings from mid-20th-century studies (Schnepp and Schnepp 1971; Rushton 1951). This model offers a more physiologically valid and statistically sound basis for interpreting g-ratio data.

Another critical issue in the g-ratio literature is the misapplication of regression slopes in scatterplots as indicators of biological significance. As reported (Gow 2025), interpretations drawn from those slopes can misrepresent true biological relationships, especially since g-ratios under healthy conditions tend to remain relatively stable across axon diameters within a given tract.

Furthermore, the same authors emphasized that many reported correlations are subject to statistical artifacts, resulting from nonuniform data distributions, inappropriate averaging of pseudo-replicate data, and failure to account for the hierarchical structure of biological measurements. These methodological flaws not only distort the interpretation of g-ratio data but can undermine the reliability and reproducibility of comparative analyses across studies.

Minor variations in g-ratio can also be overinterpreted regarding their functional impact. For instance, classical models of axonal conduction predict that increasing the g-ratio from 0.75 to 0.85 in a 1 µm axon results in only about a 5–10% reduction in conduction velocity (Waxman and Swadlow 1977; Rushton 1951). This highlights the importance of interpreting g-ratio changes within a physiological context in different axon populations. Thus, there is a need for standardized data handling practices, such as binning and clustering, which are essential for conducting biologically meaningful analyses. Binning involves grouping axons into predefined diameter intervals and calculating the average g-ratio values within each bin. This method helps stabilize variance across the dataset, thereby reduces heteroscedasticity, facilitates the detection of biologically meaningful patterns, and highlights whether specific subpopulations (e.g., small vs. large axons) are differentially affected by diseases or treatment conditions. In contrast, clustering categorizes data based on biologically relevant factors such as animal ID, experimental group, or anatomical region. This approach preserves the nested structure of the data, mitigates pseudoreplication, and enables proper statistical modeling using ANOVA or mixed-effects methods (Lazic 2010; Chomiak and Hu 2009; Gow 2025).

To overcome these longstanding challenges in g-ratio analysis, we developed MyeliMetric, a toolbox that automates and standardizes post-segmentation processing of axon and myelin data. Grounded in the axomyelin unit model, MyeliMetric integrates biologically informed data cleaning, accurate g-ratio computation, and systematic binning. By minimizing analytical bias and enhancing reproducibility, this tool provides a robust and validated framework for high-confidence g-ratio assessment across diverse experimental models and pathological conditions.

## Materials and Methods

### Python Ecosystem

MyeliMetric is developed in Python 3.7, leveraging a suite of well-established libraries for data processing, statistical analysis, and visualization. Core dependencies include Pandas for tabular data manipulation (McKinney 2010), NumPy for numerical computations (Harris et al. 2020), and SciPy for statistical tests such as the Shapiro-Wilk normality test and confidence interval estimation (Virtanen et al. 2020). For data visualization, Matplotlib (Hunter 2007) and Seaborn (Waskom 2021) are employed to generate histograms and scatter plots. Furthermore, OpenPyXL and xlrd facilitate reading and writing of Excel files, ensuring compatibility with common spreadsheet formats. All dependencies used in the development of MyeliMetric toolbox are summarized in **Table 1**.

**Table 1:** Required Python Libraries for Running the MyeliMetric Toolbox.

| Package | Purpose |
| --- | --- |
| tk | GUI framework for building the application window |
| pandas | Data handling and manipulation (e.g., Excel I/O) |
| numpy | Numerical operations |
| matplotlib | Plotting visualizations (e.g., histograms, scatter plots) |
| scipy | Scientific computations/statistics (used in analysis modules) |
| openpyxl | Read/write Excel files with pandas |
| pillow | Image loading for GUI background |
| diptest | Test for multimodality in a univariate distribution |
| xlsxwriter | Alternative engine for writing Excel files |
| statsmodels | It provides classes and functions for the estimation of statistical models |

MyeliMetric can be launched directly via the “main.py” script using any standard Python 3.7+ environment. A script named “install_dependencies.py” is provided in the script folder to streamline setup by automatically installing the required Python packages. For users without a Python installation, a portable Windows version with all dependencies pre-configured is also available as a download from the repository.

### Modular Architecture and Data Handling

MyeliMetric is designed with a modular, offline-first architecture to ensure robustness when handling large datasets and to securely manage sensitive experimental data. This modular user interface (UI) layout (**Figure 1**) separates each functional component: data import, cleaning, computation, analysis, comparison, and visualization, allowing them to operate independently while functioning cohesively. This structure enhances maintainability, simplifies debugging, and supports future expansion or integration with other analysis pipelines. Additionally, the modularity fosters transparency by clearly delineating the role and boundaries of each processing step, making the tool accessible to both novice and experienced users.

**Figure 1.**
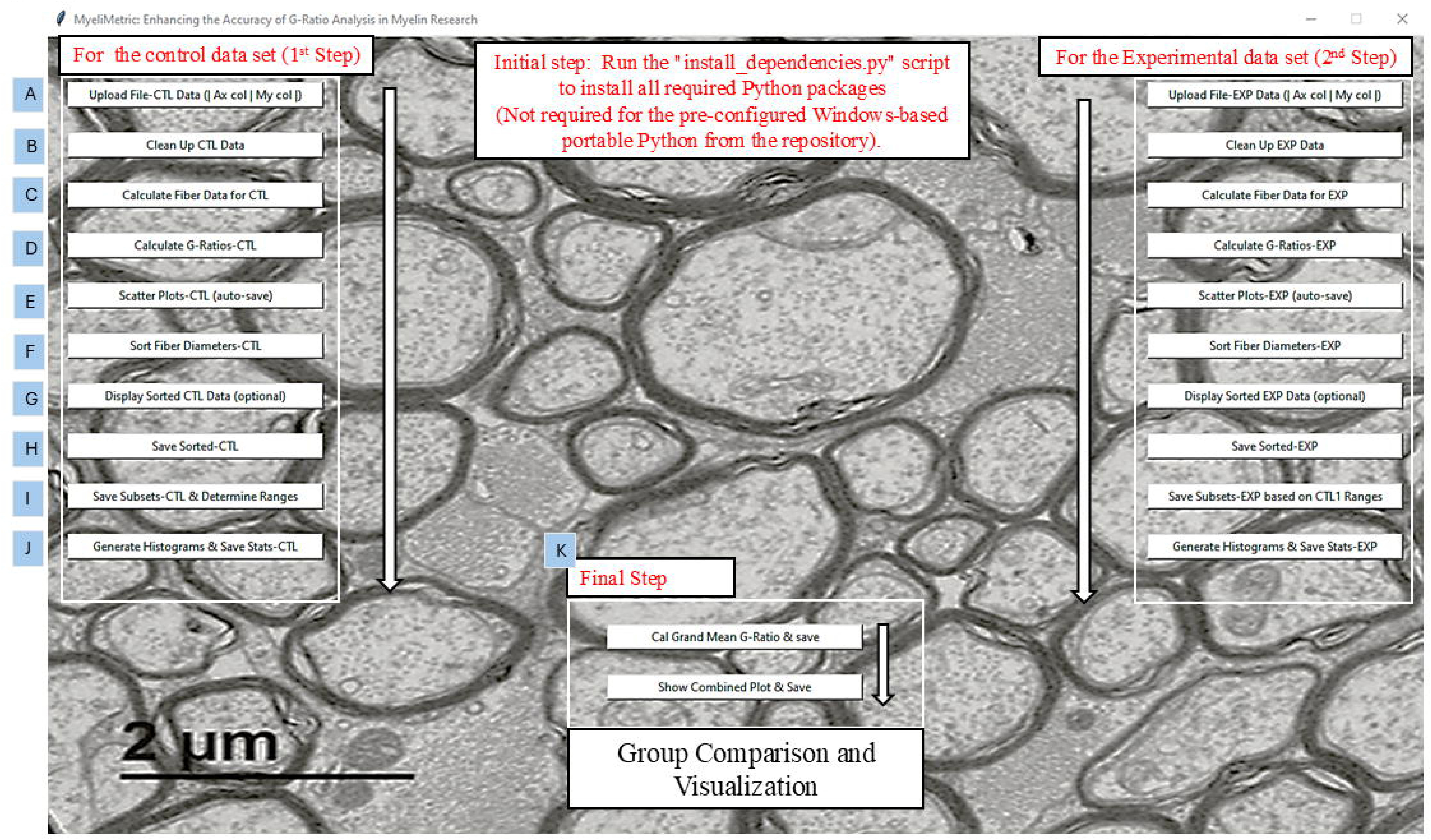
Modular graphical user interface (GUI) of the MyeliMetric toolbox. A screenshot of the opening interface of the MyeliMetric software application. The modular layout supports a stepwise workflow, beginning with dependency installation. The interface is organized into three sequential sections. Users first process the control group (left panel), which includes: (A) data upload, (B) data cleaning, (C) fiber diameter calculation, (D) g-ratio computation, (E) scatter plot generation, (F) data sorting, (G) data display, (H) output saving, (I) subset definition and reference range determination, and (J) histogram generation with statistical annotations. The experimental dataset (right panel) follows the same workflow. In the final step (K, bottom panel), users can calculate the grand mean g-ratio across groups and generate integrated visualizations, enabling comprehensive comparisons of treatment effects on myelin integrity.

The toolbox begins with the IO Handler module, which reads .xls input files, checks for required column headers (e.g., SampleName_Ax for axon diameter and SampleName_My for myelin thickness), and organizes the contents into internal pandas.DataFrame objects for downstream analysis in Python. For each sample, the column names must share an identical prefix (SampleName) to ensure the correct pairing of axon and myelin data. These input files should contain numerical values in float64 format, representing micrometer-scale measurements. If myelin thickness is provided as a one-sided radial measurement, it must be multiplied by two before analysis. Example input files conforming to this structure are provided in the repository to facilitate correct formatting and usage.

### Analytical Rationale and Workflow

To ensure biologically meaningful g-ratio calculations, in addition to removing missing entries, data cleaning thresholds are applied to exclude axon diameters smaller than 0.15 µm and myelin thicknesses below 0.03 µm. These cutoffs are supported by multiple ultrastructural studies. In 1980, it was reported that axons smaller than 0.15 µm are rare and may lack functional significance (Sturrock 1980), while Hildebrand and Hahn noted that extremely thin myelin sheaths (<0.03 µm) are unlikely to provide effective insulation (Hildebrand and Hahn 1978). More recently, it has been demonstrated that in the central nervous system, oligodendrocytes typically myelinate axons larger than 0.3 µm, reinforcing the existence of a lower functional threshold (Arancibia-Cárcamo et al. 2017). Axons smaller than ∼0.2 µm are frequently unmyelinated or inconsistently myelinated, making g-ratio estimates in this range biologically unreliable.

To facilitate stratified analysis of axonal myelination profiles, MyeliMetric implements a binning strategy that partitions axons into six predefined diameter-based subgroups. This framework accounts for the inherent heterogeneity in axon calibers observed across the central nervous system, where axon diameter is a key determinant of myelin sheath thickness and conduction velocity. Grouping axons into discrete size categories enables quantitative comparisons that can uncover diameter-specific patterns of myelin integrity, which may be obscured in bulk or non-stratified analyses. This approach is biologically justified by foundational studies demonstrating that larger axons exhibit greater reliance on myelin for efficient signal conduction and are differentially susceptible to demyelination (Hildebrand and Hahn 1978; Waxman and Bennett 1972). The use of six bins reflects an optimal compromise between analytical resolution and statistical power. Fewer bins (e.g., 2-3) may oversimplify the data and obscure subgroup-specific trends, while more bins (e.g., 10 or more) may result in sparse subgroups, reducing the robustness of statistical comparisons. Theoretically, six bins provide sufficient granularity for interpretability while maintaining adequate sample sizes within each category (Gow et al. 2025; Gow 2025). This binning strategy aligns with the broader analytical approaches used in previous ultrastructural and MRI-based g-ratio studies, which emphasize the importance of examining g-ratio variation as a function of axon caliber (Stikov et al. 2015; Arancibia-Cárcamo et al. 2017). The range boundaries for the six subgroups are derived from the distribution of axon diameters in the control dataset, which provides a representative baseline for biologically plausible values. These bin ranges are then applied consistently across all experimental groups to support standardized comparisons and maintain interpretive consistency throughout the analysis.

### Simplified Pseudocode for the Workflow Single-Group G-ratio Analysis

BEGIN
LOAD input Excel files (e.g., control or experimental group)
VERIFY required columns (e.g., SampleName_Ax and SampleName_My) CLEAN data:
REMOVE entries with missing values APPLY thresholds:

– EXCLUDE axon diameter < 0.15 µm
– EXCLUDE myelin thickness < 0.03 µm LOG all exclusions and modifications
CALCULATE fiber diameter = axon diameter + myelin thickness CALCULATE g-ratio = axon diameter/fiber diameter
BIN axons into 6 predefined diameter categories:

– Bin ranges derived from the control dataset (CTL1) distribution
– APPLY the same binning ranges to all groups for consistency FOR each bin:
CALCULATE mean, median, and standard deviation PERFORM normality test (i.e., Shapiro-Wilk) GENERATE histograms and scatter plots
RETURN summary statistics and visual outputs
END

### G-ratio Comparison

BEGIN
LOAD data for control and experimental groups
VALIDATE presence of required columns: axon_diameter, myelin_thickness CLEAN both datasets (remove missing or invalid entries)
IF both control AND experimental datasets are available:
FOR each matching bin:
PERFORM statistical comparison (i.e., the 2-way ANOVA)
\# Comparative statistical analysis can be performed on the output as needed, depending on the experimental design.
GENERATE comparative plots:

– Histograms of g-ratios
– Scatter plots of g-ratio vs axon diameter
SAVE outputs (plots, mean ± SD, and cleaned data) to the specified output directory END

To enhance transparency and reproducibility, the modular architecture of MyeliMetric is detailed in **Figure 2**, which outlines the scripts, their core functions, operational formulas, and specific roles within the analysis pipeline. This structured breakdown enables users to understand the internal logic of the software and facilitates the selective execution of individual components as needed.

**Figure 2.**
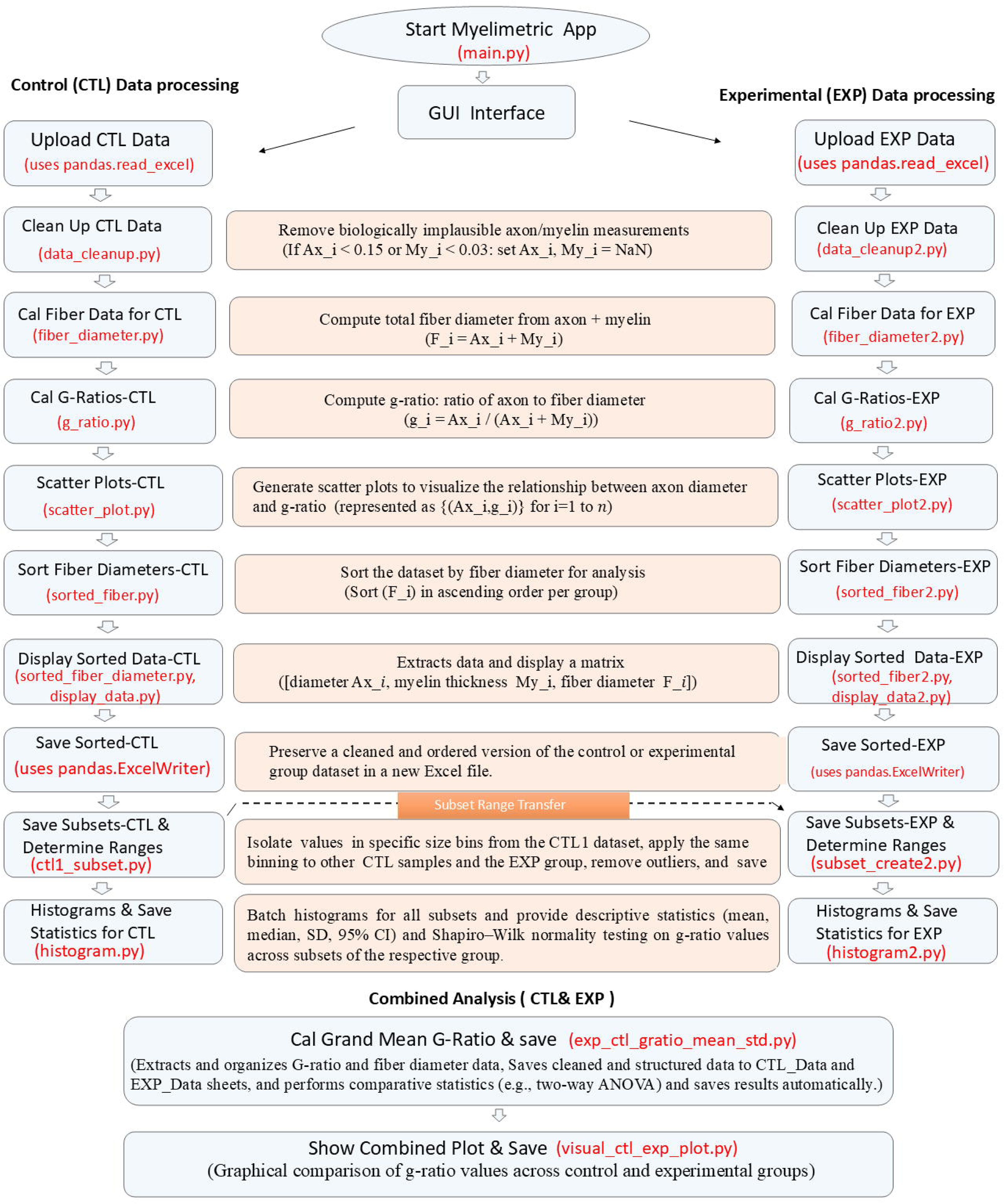
Architecture of the MyeliMetric toolbox. The MyeliMetric pipeline consists of distinct but interconnected scripts organized into functional modules for processing control (CTL) and experimental (EXP) datasets. Each module performs a specific analytical task, including data cleaning, fiber diameter calculation, g-ratio computation, scatter plotting, data sorting, and statistical evaluation. Buttons in the GUI are annotated with their corresponding implementing scripts, providing a transparent and traceable workflow.

### Evaluation Plan

MyeliMetric was validated through a three-tier process to ensure reliability and performance. In the first tier, unit tests were performed on custom-developed modules that are not part of any existing Python libraries. These tests were used to simulate edge cases such as missing columns, extreme values, and inconsistent data formats. All operations were systematically logged to ensure full traceability and reproducibility of the results. In the second tier, a large synthetic dataset was generated by mimicking the statistical properties of real experimental data, allowing performance benchmarking under realistic conditions. In the third tier, the software was applied to published demyelination datasets to assess its ability to reproduce known g-ratio patterns. All unit test scripts are provided in the repository to ensure reproducibility and transparency.

### Synthetic Data Generation Reflecting Real Experimental Properties

To evaluate the accuracy and robustness of g-ratio computations under controlled conditions, a synthetic dataset was generated that mimics the statistical properties of real axon and myelin measurements observed in our control samples. This approach ensures that the data structure and variability resemble biologically plausible distributions while allowing us to define and trace ground-truth g-ratio values. Descriptive statistics were first calculated from real measurements of axon diameter and myelin thickness extracted from five biological replicates of control animals (Dupree et al. 2022). The interquartile range (IQR) for axon diameter was approximately 0.3 - 1.2 µm, and the typical myelin thickness ranged between 0.12 and 0.44 µm. The estimated mean g-ratio from real data was approximately 0.6, aligning with expected physiological ranges in the CNS tissue.

Using these empirical observations, we generated 1,000 synthetic axons, myelin, and fiber diameter triplets. Axon diameters were randomly sampled from a uniform distribution spanning 0.3 - 1.2 µm, consistent with the IQR of the example empirical dataset. Myelin thicknesses were computed using a base scaling factor (base ratio = 0.4) applied to each axon diameter, modulated by random biological variability to simulate natural noise. This was implemented as: Myelin = Axon × base ratio × ε, where ε ∼ U(0.7, 1.3). This random multiplier ε simulates variation in oligodendrocyte wrapping efficiency and biological dispersion. Fiber diameters were calculated as: Fiber = Axon + (2 × Myelin), ensuring a realistic structure where myelin is bilaterally wrapped around the axon. G-ratio (g) and mean g-ratio (ḡ) values were derived using the canonical formula:

1. g-ratio (g) = Inner diameter of the myelin sheath (axon diameter) / Outer diameter of the myelin sheath (axon + myelin)
2. mean g-ratio (ḡ) = (1/n) × Σ [aᵢ/(aᵢ + 2mᵢ)], for i = 1 to n, where: n is the total number of axons, aᵢ is the diameter of the i-th axon, mᵢ is the myelin thickness of the i-th axon.

## Results

### Synthetic Data Validation

Unit tests were conducted on all custom-developed modules to validate their functional correctness and robustness. The results confirmed that each function executed as intended, produced accurate and consistent outputs, and effectively handled atypical input scenarios, including edge cases. No functional errors or unexpected behaviors were detected during testing. A comprehensive summary of the tested modules, test scope, outcomes, and associated external dependencies is presented in **Table 2**.

**Table 2:** Verification of Core Functions in the MyeliMetric Toolbox via Unit Testing (External Module Dependencies)

| Unit test script | Module to be tested | Tested | Status |
| --- | --- | --- | --- |
| test_data_cleanup.py | data_cleanup.py<br>data_cleanup2.py | Verifies filtering based on thresholds (axon < 0.15, myelin < 0.03) and realignment of non-NaN values | **Pass** |
| test_subset.py | ctl1_subset.py | Functions related to subset creation and range saving<br>• Test subset creation and min/max validation<br>• Test filtering and correct sheet naming<br>• Test saving range data and removing empty sheets | **Pass** |
| test_g_ratio.py | g_ratio.py<br>g_ratio2.py | Correctness of g-ratio values ($Ax / (Ax + My)$, rounded to 2 decimals) and verifies input columns are present and expected output columns are created | **Pass** |

In a synthetic dataset comprising 1,000 axons, g-ratio values were randomly generated from a distribution with a mean closely approximating 0.6 (μ=0.59, σ²=0.002), effectively capturing the natural variability observed in healthy myelinated fibers (**Figure 3A**). Importantly, we observed a strong positive correlation between axon diameter and fiber diameter (Pearson’s r ≈ 0.9). It supports the phenomenon that as axon diameter increases, myelination typically increases to preserve the optimal g-ratio, which ensures efficient nerve conduction. However, fiber diameter does not correlate with g-ratio highlighting the expected independence of these two variables (**Figure 3B**). When the data were sorted into six subsets based on fiber diameter, the observed trend demonstrated that the binning approach preserved the natural spread of fiber sizes while effectively stratifying the dataset for downstream analysis of the grand mean g-ratio from each subset (**Figure 3C**). The mean g-ratio remained consistent at approximately 0.6 across all subsets, aligning with the principle that g-ratio should be stable within a homogeneous nerve fiber population. The final grand mean g-ratio was 0.59, matching the manually calculated value from the entire (n=1000) dataset (**Figure 3D**).

**Figure 3.**
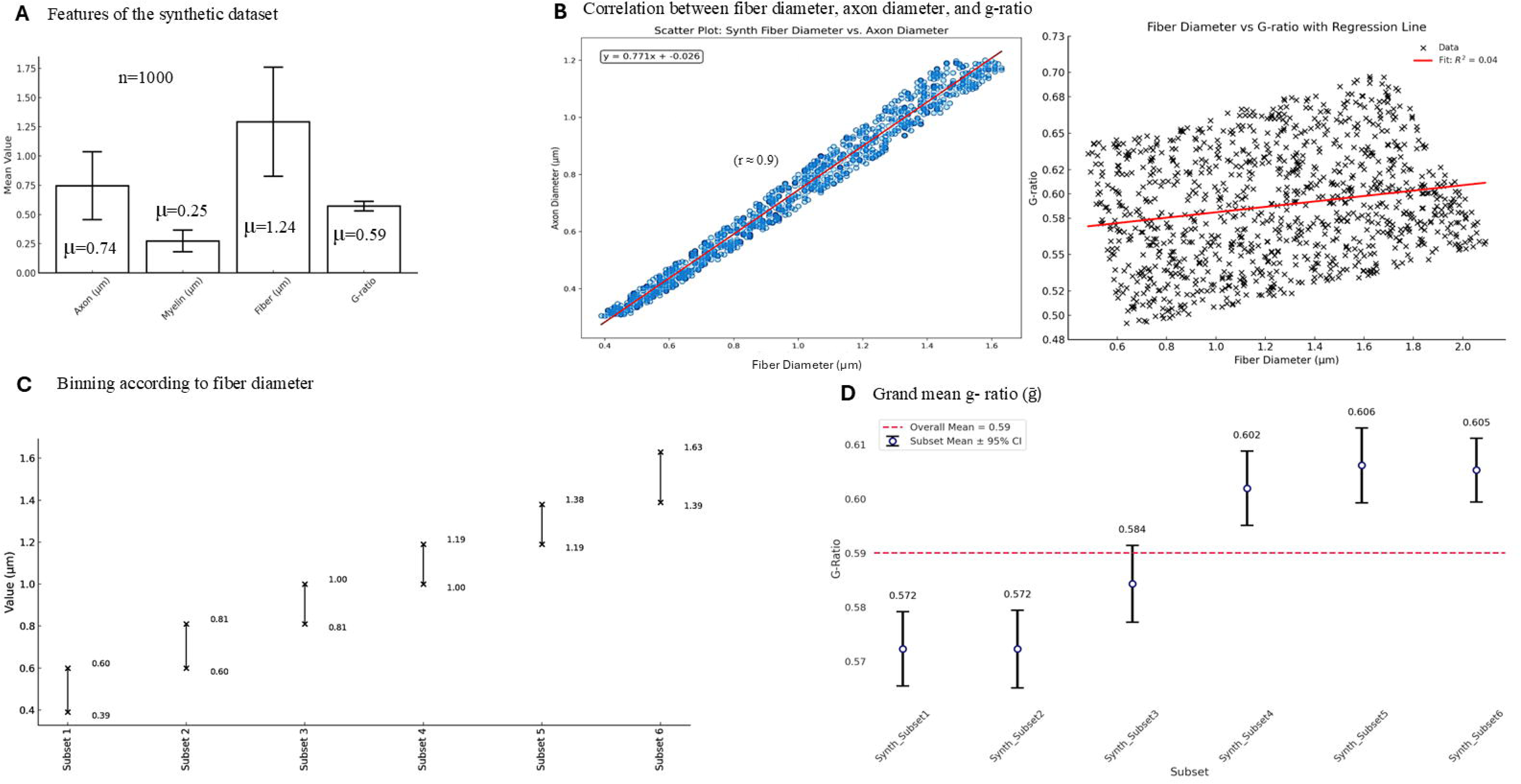
Validation of g-ratio distribution and correlation patterns using a synthetic dataset. (A) A synthetic dataset of 1,000 axons was generated with g-ratio values drawn from a normal distribution centered at 0.6 (μ = 0.59, σ² = 0.002), replicating natural variability in healthy myelinated fibers. (B) The left panel shows a strong positive correlation between axon diameter and fiber diameter (Pearson’s r ≈ 0.9), supporting the concept that increased axon size is typically accompanied by greater myelination to maintain optimal conduction. The right panel shows no significant correlation between fiber diameter and g-ratio, confirming the expected independence of these variables. (C) The dataset was divided into six subsets based on fiber diameter. This binning approach preserved the distribution of fiber sizes and enabled stratified analysis. (D) The mean g-ratio remained consistent (∼0.6) across all subsets, validating the homogeneity and biological plausibility of the synthetic model. The grand mean g-ratio (red line) was 0.59, identical to the mean of the full dataset (n = 1,000).

### Validation Against Published Data

To test MyeliMetric against real-world experimental data, we re-analyzed a published dataset from a study examining cuprizone-induced demyelination in mice with and without Lanthionine Ketimine Ethyl Ester (LKE) supplementation (Dupree et al. 2022). The grand mean g-ratio values (ḡ) calculated by MyeliMetric closely match the trends reported in the original study, confirming the consistency and reliability of our tool in reproducing biologically meaningful results from published datasets (**Figure 4A-C**). Minute deviations were observed, as expected, mainly due to differences in outlier exclusion, binning strategies, and data handling protocols used by MyeliMetric, which implements stricter filtering criteria for physiological plausibility.

**Figure 4.**
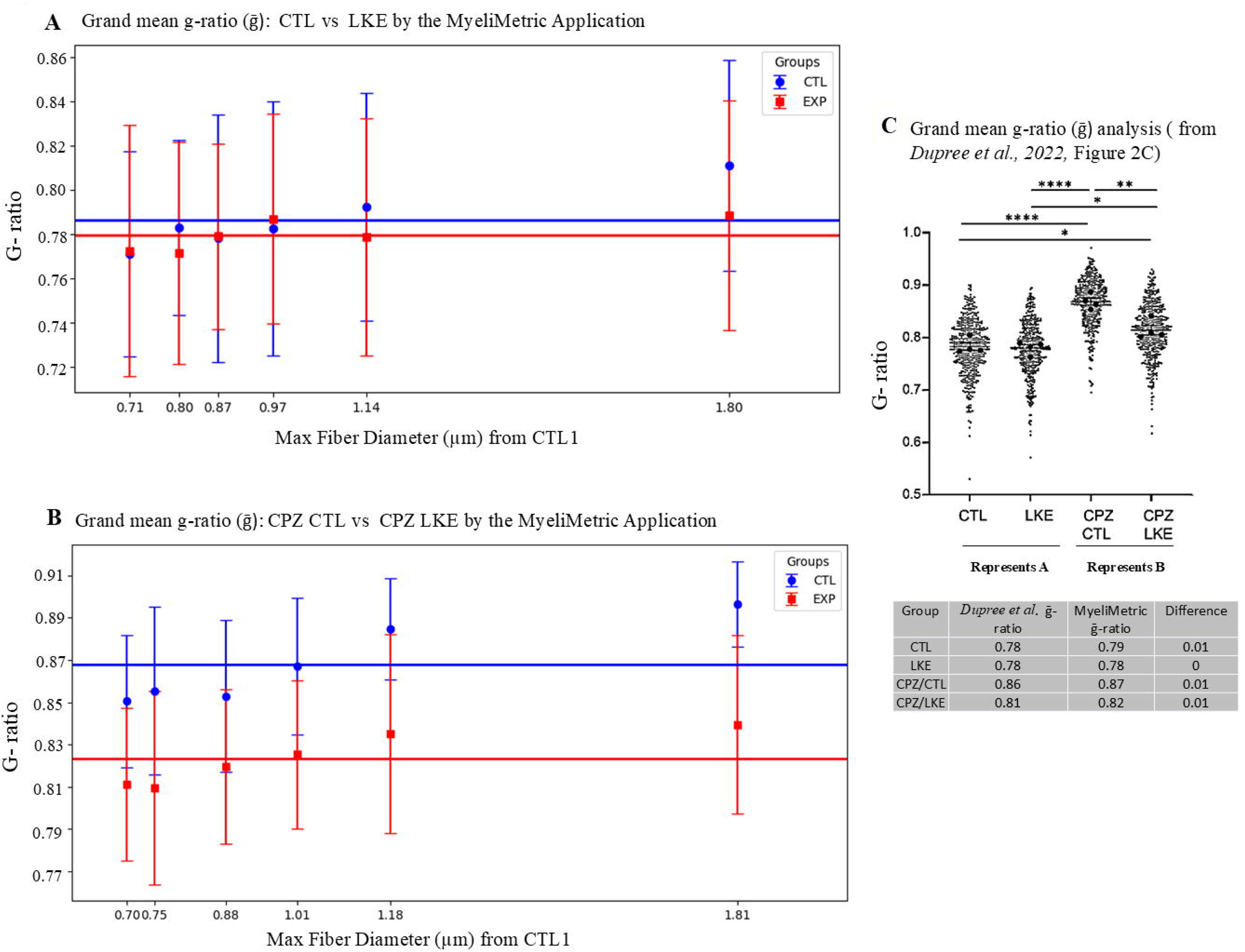
Validation of MyeliMetric using a published dataset on cuprizone-induced demyelination and LKE treatment. Grand mean g-ratio (ḡ) values were calculated using MyeliMetric for (A) Control (CTL) and LKE-treated groups and (B) cuprizone-treated mice followed by either control (CPZ/CTL) or LKE (CPZ/LKE) chow. (C) The computed grand mean g-ratio values were compared with those reported by Dupree et al. (2022). The strong concordance validates the tool’s reliability and accuracy in analyzing real-world datasets.

Importantly, MyeliMetric introduced enhanced analytical granularity by segmenting axons into diameter-based bins, revealing fiber-size-specific responses to LKE treatment (**Figure 4A,B**). For instance, while the original study reported a significant reduction in ḡ-ratio following LKE administration after two weeks of remyelination (CPZ/LKE vs. CPZ/CTL), a 2-way ANOVA analysis on data generated by MyeliMetric demonstrated that the Group × Subset (fiber diameter bins) interaction approached but did not reach statistical significance (F (5, 994) = 1.99, p = 0.078).

This implies that while both group and subset independently influence g-ratio values, there is no strong evidence that the effect of group differs systematically across fiber diameter subsets. Additionally, Shapiro–Wilk normality testing in subgroups revealed deviations from Gaussian distribution in some bins. This statistical profiling (provided in spreadsheet) gives a high-level overview of data structure and variability (**Figure 5A**). MyeliMetric also generated scatter plots of fiber diameter versus axon diameter for individual samples, alongside subset-specific histograms for each subject (**Figure 5B**), providing a detailed visualization of axon–myelin relationships. These plots facilitate a deeper understanding of the distributional properties within and across fiber diameter subsets, thereby enhancing the interpretability at both the individual and subset (bin) levels.

**Figure 5.**
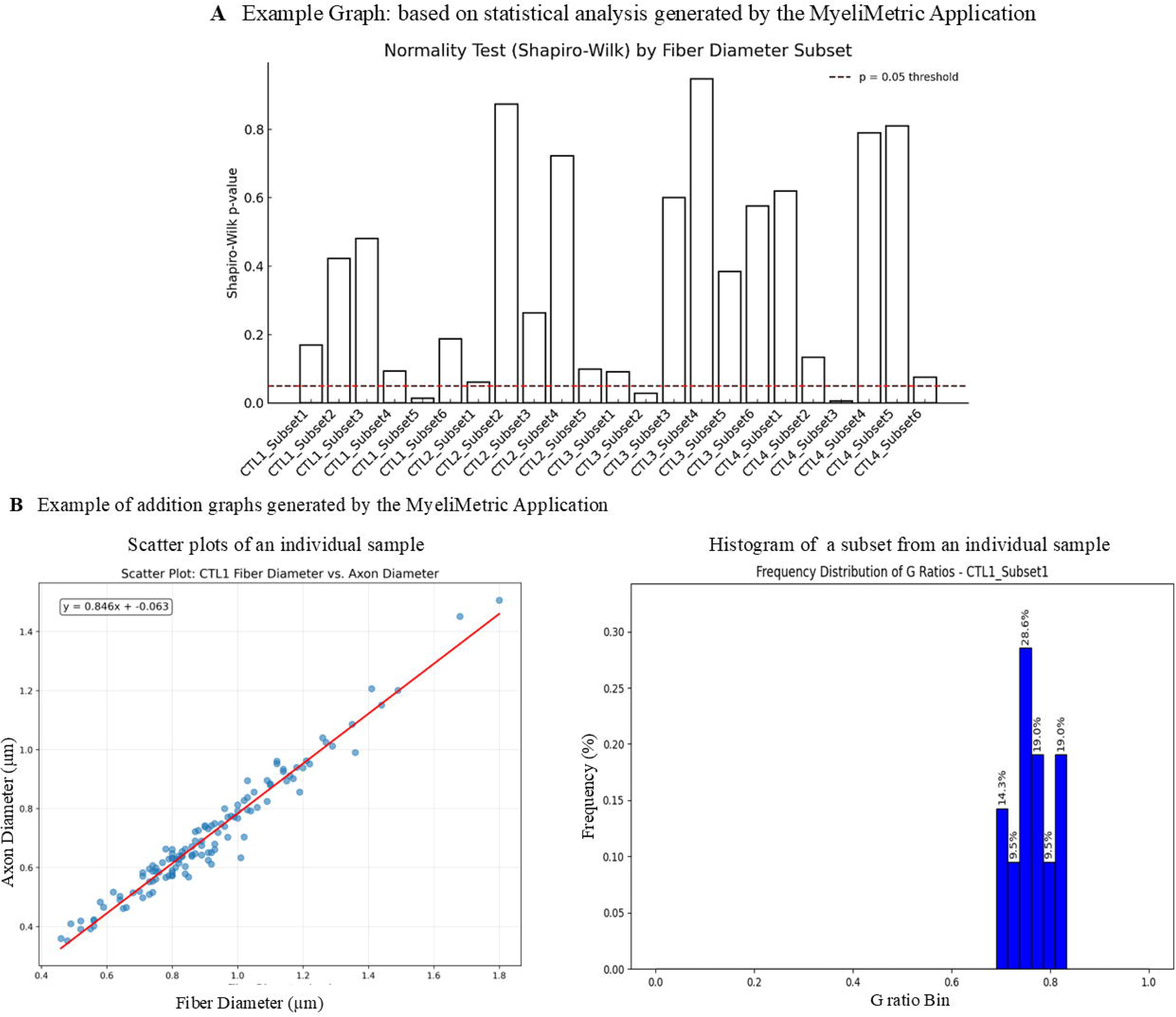
Statistical and visual profiling of diameter-binned g-ratio data generated by MyeliMetric. (A) Shapiro–Wilk normality testing across fiber diameter subsets identified deviations from Gaussian distribution in three bins (p < 0.05, red line), highlighting data heterogeneity. (B) MyeliMetric-generated scatter plots of axon diameter versus fiber diameter, along with subset-specific histograms for individual subjects, provide detailed visualizations of axon–myelin relationships and enhance interpretability at both individual and group levels.

## Discussion

MyeliMetric advances the current landscape of g-ratio analysis; it is a purpose-built Python toolbox that enhances the post-segmentation analysis of axon–myelin structures by implementing biologically grounded and statistically rigorous workflows. Rather than relying on fragmented or ad hoc approaches, MyeliMetric offers an integrated solution that improves the reliability of g-ratio estimation through standardized data processing and contextual validation. Its performance demonstrated across both synthetic and experimental datasets highlights its utility in uncovering subtle, fiber-size-dependent patterns in myelin architecture, an advancement over conventional methods that often overlook such granularity.

Enhanced resolution in g-ratio analysis is particularly important in demyelinating conditions such as multiple sclerosis (MS), where remyelination dynamics can vary significantly across axon calibers. Previous studies have shown that large-caliber axons tend to exhibit greater spontaneous remyelination capacity during recovery (Franklin and Goldman 2015), highlighting the importance of size-specific assessment. This pattern reinforces the need for granular post-segmentation analysis to accurately evaluate treatment effects. Notably, such selective remyelination aligns with findings of enriched populations of specific remyelinating oligodendrocytes, such as ermin-expressing cells within active repair zones of MS lesions (Ahmad et al. 2021). These observations collectively emphasize that therapeutic responses are not uniformly distributed across all axons, and tools like MyeliMetric are essential for detecting and interpreting such biologically relevant heterogeneity.

Given that control datasets serve as the physiological baseline, it is both reasonable and necessary to apply biologically informed filters during analysis. The inclusion of implausible values, such as those resulting from segmentation artifacts or imaging inconsistencies that can distort statistical measures and lead to misleading subgroup classifications. In contrast, experimental datasets may legitimately contain atypical values that reflect pathological or regenerative changes, including severely demyelinated axons or unusually thin myelin sheaths. To account for this distinction, MyeliMetric incorporates a flexible filtering system that allows users to enable or bypass the data cleanup module based on the biological context. Its modular design supports this adaptability while maintaining analytical integrity. Furthermore, all excluded data are automatically logged in a separate output file, promoting transparency and enabling users to review, audit, or reprocess the filtered entries as needed.

Another crucial advantage in analysis with the MyeliMetric application is its integration of biological plausibility checks. The software flags unexpected correlations between fiber diameter and g-ratio, prompting users to investigate potential methodological artifacts such as segmentation bias, inconsistent imaging resolution, or sampling error (Chomiak and Hu 2009). Additionally, g-ratio values falling outside of the expected physiological range (typically <0.5 or >0.9) are automatically removed as potential outliers. These built-in quality control features are essential for ensuring scientific rigor and reproducibility, particularly in multicenter or longitudinal studies where methodological variability can be a significant confounding factor.

Simulated datasets modeled after empirical axon and myelin measurements offered a controlled environment to evaluate the reliability of MyeliMetric under physiologically realistic conditions. The tool consistently returned g-ratio values within the expected range while filtering out entries that lacked biological plausibility. This performance is particularly important given the narrow window within which g-ratios are considered physiologically normal.

A re-analysis of experimental data from a cuprizone-induced demyelination model (Dupree et al. 2022) further highlights the utility of MyeliMetric. While the original study reported a significant reduction in g-ratio following LKE treatment, our stratified analysis revealed a more nuanced effect, with group × subset interactions (based on fiber diameter bins) approaching statistical significance. This finding underscores the value of bin-wise comparisons and mixed-model approaches, particularly in studies with moderate sample sizes or heterogeneous axonal populations (Lazic 2010; Gow et al. 2025). Additionally, the identification of non-Gaussian distributions in several diameter bins reinforces the need to incorporate normality testing as a routine step in g-ratio analyses, an aspect often neglected in previous work.

While tools such as AxonDeepSeg (Zaimi et al. 2018) and MyelTracer (Kaiser et al. 2021) provide high-quality segmentation of axons and myelin from microscopy images, they do not offer integrated post-processing pipelines for g-ratio computation, outlier filtering, or biologically contextualized analysis. MyeliMetric fills this critical gap by providing a modular, transparent framework that enhances reproducibility and reduces user-induced variability. Designed to complement existing segmentation platforms, MyeliMetric streamlines post-segmentation workflows and supports downstream export to statistical environments such as R, Python’s statsmodels, or machine learning frameworks. This interoperability is especially important as neuroscience increasingly embraces integrated, multi-modal analyses. By enabling high-confidence morphological quantification, MyeliMetric supports the alignment of imaging-based findings with transcriptomic, proteomic, and behavioral data, facilitating deeper insights into both systems neuroscience and translational research.

Despite its strengths, MyeliMetric has several limitations. First, the toolbox relies on the accuracy of upstream segmentation and cannot validate or correct errors introduced during image preprocessing or axon/myelin labeling (Zaimi et al. 2018). As a result, it may not adequately account for irregularly shaped or obliquely sectioned axons, which can distort area-to-diameter conversions. Second, while the tool is optimized for high-resolution electron microscopy data, its performance on lower-resolution or noisy datasets, such as those generated by light microscopy requires further validation. Additionally, the current version supports only .xlsx file formats, limiting compatibility with alternative data structures commonly used in large-scale or automated pipelines. At present, advanced statistical analyses must be performed externally, as they are not integrated into the software. Future iterations may include 3D volumetric support, machine learning–based outlier detection, expanded file format compatibility, and embedded statistical comparison modules to enhance both scalability and user accessibility.

In summary, MyeliMetric addresses a critical methodological need in neurobiological research by offering a validated, reproducible, and user-friendly pipeline for g-ratio analysis. It represents a significant advancement in g-ratio quantification by providing a biologically grounded, quality-controlled, and statistically robust analysis pipeline. Its ability to resolve fiber-size-specific patterns, detect artifacts, and support reproducible workflows addresses long-standing challenges in axon–myelin research. As experimental neuroscience continues to evolve toward higher-resolution, multimodal investigations, MyeliMetric offers a foundational tool for high-confidence morphometric analysis in both basic and translational myelin research.

## Supporting information

Sample_DataSets

